# The effect of sex and underlying disease on the genetic association of QT interval and sudden cardiac death

**DOI:** 10.1101/664300

**Authors:** Rebecca N. Mitchell, Foram N. Ashar, Marjo-Riitta Jarvelin, Philippe Froguel, Nona Sotoodehnia, Jennifer A. Brody, Sylvain Sebert, Heikki Huikuri, John Rioux, Philippe Goyette, Charles E. Newcomb, M. Juhani Junttila, Dan E. Arking

**Author notes:** ^#^**Corresponding Author:** Dan E. Arking, PhD, Johns Hopkins University School of Medicine, 733 N. Broadway, Miller Research Building, Room 459, Baltimore, MD 21205.

## Abstract

**Background:** Sudden cardiac death (SCD) accounts for ~300,000 deaths annually in the US. Men have a higher risk of SCD and are more likely to have underlying coronary artery disease (CAD) than women. In contrast, women are more likely to have arrhythmic events in the setting of inherited or acquired QT prolongation. Moreover, there is evidence of sex differences in the underlying genetics of QT interval duration. Using sex- and CAD-stratified analyses, we assess differences in genetic association between prolonged QT interval and SCD risk.

**Methods:** We examined 2,282 SCD subjects with autopsy-confirmed underlying disease from the Fingesture cohort and 3,561 Finnish controls. The SCD subjects were stratified by underlying disease (ischemic vs. non-ischemic) and by sex. We used logistic regression to test for association between the top QT interval associated SNP, rs12143842 (in the *NOS1AP* locus), and SCD risk. We also performed Mendelian randomization to test for causal association of QT interval in the various subgroups.

**Results:** Female SCD victims with underlying non-ischemic disease had the strongest association between rs12143842 and SCD risk (OR=1.37; 95% CI, 1.07-1.75) and the strongest causal association, established using Mendelian randomization, between prolonged QT interval and SCD (OR in SCD risk per SD increase in QT, 3.60; 95% CI, 1.22-10.49). Ischemic SCD victims, irrespective of sex, did not show an association between rs12143842 and SCD risk or a causal association for QT interval.

**Conclusions:** This study provides evidence that the causal effect of QT prolongation on SCD risk differs by sex and underlying disease.

## INTRODUCTION

Sudden cardiac death (SCD) is among the leading causes of death in the United States, affecting approximately 300,000 individuals annually.^1^ SCD occurs as a result of multiple underlying disease pathologies, including heart diseases such as coronary artery disease (CAD) and cardiomyopathies, as well as primary electrical defects.^2^ Men have a higher risk of SCD than women^3,4^, and furthermore, the underlying cardiac pathology differs between the sexes. CAD, the common underlying cause of SCD, is more common in men than women. By contrast, non-ischemic pathology, such as primary myocardial fibrosis, valvular heart disease, and arrhythmogenic right ventricular cardiomyopathy, occurs more commonly in women with SCD compared to men with SCD.^5–7^ SCD is often the first manifestation of heart disease, particularly for women; several studies have found that women are less likely than men to have a prior history of known cardiac disease.^4,8^ It has been hypothesized that SCD is a much more heterogeneous condition in women, potentially due to the different underlying diseases, leading to differences in the associated risk factors.

Prolonged QT interval, a measure of ventricular repolarization, has been previously established as a risk factor for SCD,^9,10^ and recent studies using Mendelian randomization have demonstrated that this risk factor is causal.^11^ Women, on average, exhibit longer QT intervals than men in the general population once puberty is reached.^12,13^ In addition, a previous study found that the increase in risk for overall cardiac death associated with prolonged QT interval was more pronounced in women.^14^ Women also have higher risk of arrhythmic events than men in the setting of inherited or acquired (drug-induced) QT prolongation.^15^ Based on the sex differences in QT interval in the general population and its association with overall cardiac mortality, we hypothesize that the risk of SCD associated with prolonged QT interval could differ by sex. Likewise, we also hypothesize that QT interval could differentially affect SCD risk depending on the underlying pathology (e.g. ischemic vs. non-ischemic disease).

Previous studies have shown that ~34% of QT interval variation is heritable^16,17^. In addition, recent research indicates that ~21% of variation can be explained by common autosomal SNPs found genome-wide, including SNPs in genes such as *KCNQ1*, *KCNH2*, *SCN5A* and *NOS1AP*.^18^ The top SNP from the most recent QT interval genome-wide association study (GWAS) was the *NOS1AP* locus SNP rs12143842, which increased QT interval by 3.50 ms per T-allele (p-value=1×10^−213^)^19^ and accounts for ~1% of the variation in QT interval.^20^ This SNP has been previously associated with increased SCD risk^21,22^, and has also been found to have stronger effect on QT interval in women than men.^20^

In this study we examined a large Finnish study of post-mortem autopsy-confirmed SCD subjects to study the genetic association between QT interval and SCD risk. More specifically, we compared the association of the *NOS1AP* locus variant rs12143842 with SCD risk between subjects with underlying ischemic vs. non-ischemic disease. We also performed sex-stratified analyses within these groups to investigate any sex-specific association of the *NOS1AP* locus SNP with SCD risk. Finally, we performed Mendelian randomization to test for differences in the causal association between a previously identified causal risk factor, prolonged QT interval, and SCD in the setting of different underlying disease and/or between sexes.

## METHODS

### Samples

#### Fingesture

The Fingesture study, started in 1998, aimed to collect consecutive victims of out-of-hospital sudden death from a defined geographical area, Oulu University Hospital District in northern Finland. All victims of sudden death were autopsied at the Department of Forensic Medicine, University of Oulu, Oulu, Finland. SCD victims were defined as those with a witnessed sudden death within 6 hours of the onset of the symptoms or within 24 hours of the time that the victim was last seen alive in a normal state of health. Individuals with age at SCD event <30 years old or >80 years old were excluded from analysis.

The underlying pathologies were divided into three categories: (1) ischemic, (2) non-ischemic, and (3) other disease. The ischemic SCD victims included individuals with evidence of a coronary complication, defined as a fresh intracoronary thrombus, plaque rupture or erosion, intraplaque hemorrhage, or critical coronary stenosis (>75%) in the main coronary artery. The non-ischemic SCD victims included individuals with the following conditions: hypertrophy due to hypertension; valve disease; cardiomyopathy due to alcohol use; dilated cardiomyopathy; hypertrophic obstructive cardiomyopathy; cardiomyopathy due to obesity; arrhythmogenic right ventricular cardiomyopathy; and primary myocardial fibrosis. Further definitions of these conditions have been previously described.^5^ The “other” SCD victims included individuals with the following conditions: myocarditis, cardiac anomaly, and normal autopsy individuals (e.g. individuals with a channelopathy).

#### NFBC1966

The Northern Finland Birth Cohort (NFBC) study is the product of a project initiated in the 1960s to examine risk factors involved in pre-term birth and intrauterine growth retardation, and the consequences of these early adverse outcomes on subsequent morbidity. The NFBC1966 cohort comprised of 12,068 mothers and 12,231 children with an expected date of birth in 1966 within the province of Oulu, Finland. Our study samples consisted of DNA extracted from the blood of the offspring at their 31-year follow-up visit.

### Genotyping

Samples were genotyped for rs12143842 using five different platforms: Illumina Infinium Global Screening Array (GSA); Affymetrix Genome-wide Human SNP Array 6.0; Agena Biosciences MassARRAY; Applied Biosystems Taqman real-time PCR; and Illumina TruSeq sequencing. All genotyping and sequencing were performed according to the manufacturer’s instructions. Quality control was performed separately on each dataset before merging. Dataset and QC information is summarized in Supplementary Table 1. Overlapping samples between platforms were used to evaluate the accuracy of the genotyping (reported in Supplementary Table 1). After exclusions, the study population included 2,282 SCD victims and 3,561 Finnish controls.

### Statistical Analysis

P-values for differences in the Fingesture study characteristics were calculated using a two sample t-test for continuous variables and Pearson chi-square test for categorical variables. The genotypes for rs12143842 for all samples were merged and logistic regression was performed using R (version 3.3.3), with sex as the only covariate. The SCD cases were stratified by sex and underlying disease (ischemic, non-ischemic and other disease) to examine the SNP effects in each group. Differences between sexes were determined by incorporating an interaction term into the regression model. P-values for differences in effect sizes between the underlying disease groups were obtained from a 1-degree of freedom Wald test. Multi-dimensional scaling (MDS) using PLINKv1.9 was used for samples run on the GSA microarray (1,171 cases/761 controls) to assess potential population substructure between the Fingesture and NFBC1966 studies.

### Mendelian Randomization

Mendelian randomization uses genetic variants as instrumental variables to test for causal relationships between a trait (i.e. QT interval) and an outcome (i.e. SCD). We used a multi-SNP genetic risk score association (GRSA) model to test for causality between QT interval and SCD in our stratified datasets. The GRSA model uses 57 LD-pruned genome-wide significant SNPs from the QT interval GWAS^19^ to compare the association of these SNPs with the trait of interest (β_trait_) to the association of the SNPs with SCD (β_outcome_) using the R package ‘MendelianRandomization’.^23^ Zero-intercept inverse-weighted (IVW) linear regression is used to calculate the GRSA estimate, which is the slope of the resultant regression line, and estimates the difference in log odds of SCD risk per SD increase in QT interval. We used the HEIDI-outlier method from the ‘gsmr’ R package to detect and remove potentially pleiotropic SNPs.^24^ P-values for difference in GRSA estimates were obtained from a 1-degree of freedom Wald test.

Genome-wide SNP data is required for Mendelian randomization analyses and therefore only the Fingesture and NFBC1966 samples genotyped using the Infinium Global Screening Array (GSA) and imputed to the NHLBI Trans-Omics for Precision Medicine (TOPMed) imputation panel using the University of Michigan imputation server^25^ were used in this analysis (1,168 SCD victims and 761 Finnish controls). Logistic regression for single SNP association tests were run using FASTv2.4.^26^ We performed several stratified analyses, including by sex and underlying disease (ischemic and non-ischemic disease). There were a small number of SCD cases with other underlying disease genotyped on this array and therefore were only included in the overall analysis and sex-stratified analyses but were excluded from the underlying disease-stratified analysis and subsequent sex-stratified analyses.

## RESULTS

The SCD population is comprised of a subset of the Fingesture study of Finnish SCD subjects with autopsy-confirmed assessment of underlying heart disease in whom DNA was available at the time of this study (n=2,282). Controls were drawn from the Northern Finland Birth Cohort of 1966 (NFBC1966) and are comprised of 3,561 Finnish individuals born in 1966. Characteristics of the Fingesture study are detailed in Table 1. To assess for potential population stratification, we ran multi-dimensional scaling (MDS) on a subset of the samples with genome-wide SNP data (1,168 cases/761 controls). Plots for the top 10 MDS components, colored by SCD status, are found in **Supplementary Figure 1**. MDS component 7 was associated with SCD status after multi-test correction (*P*<0.002) (**Supplementary Table 2**), indicating the potential for confounding due to population substructure. However, combined, the top 10 components explained only 0.9% of the variance in SCD status, suggesting likely minimal impact. This minimal impact was confirmed by sensitivity analyses (described below).

**Table 1.** Fingesture study characteristics.

| **Variable** | **All**<br>**(N=2,282)** | **Men**<br>**(N=1,862)** | **Women**<br>**(N=420)** | ***P*\*** |
| --- | --- | --- | --- | --- |
| Mean age, year (SD) | 61.23 (10.71) | 60.65 (10.43) | 63.84 (11.56) | <0.001 |
| N, ischemic disease (%) | 1,478 (64.8%) | 1,245 (66.9%) | 233 (55.5%) | <0.001 |
| N, non-ischemic disease (%) | 750 (32.8%) | 579 (31.1%) | 171 (40.7%) | <0.001 |
| N, other (%) | 54 (2.4%) | 38 (2.0%) | 16 (3.8%) | 0.03 |
| BMI, kg/m<sup>2</sup> (SD) | 28.36 (6.61) | 28.16 (6.23) | 29.26 (8.10) | 0.06 |
| Heart weight, g (SD) | 493.60 (129.23) | 509.60 (127.83) | 421.40 (109.47) | <0.001 |

\**P* calculated for difference between men and women

### *NOS1AP* locus SNP analysis

Given the previously established relationship between QT interval and SCD risk, and with *NOS1AP* locus SNPs and SCD in other cohorts^10,27^, we first sought to assess the association between SCD and the *NOS1AP* locus SNP rs12143842. When analyzing all 2,282 SCD cases and 3,561 controls, the T allele of rs12143842 was significantly associated with increased SCD risk with an OR of 1.14 for each copy of the QT prolonging allele (95% CI, 1.04-1.25; *P* = 0.005). In sensitivity analyses, including the 10 top components from the MDS analysis in the model minimally increased the effect estimate (see **Supplementary Table 3**). All SNP association results are summarized in **Figure 1** and **Supplementary Table 4**.

**Figure 1.**
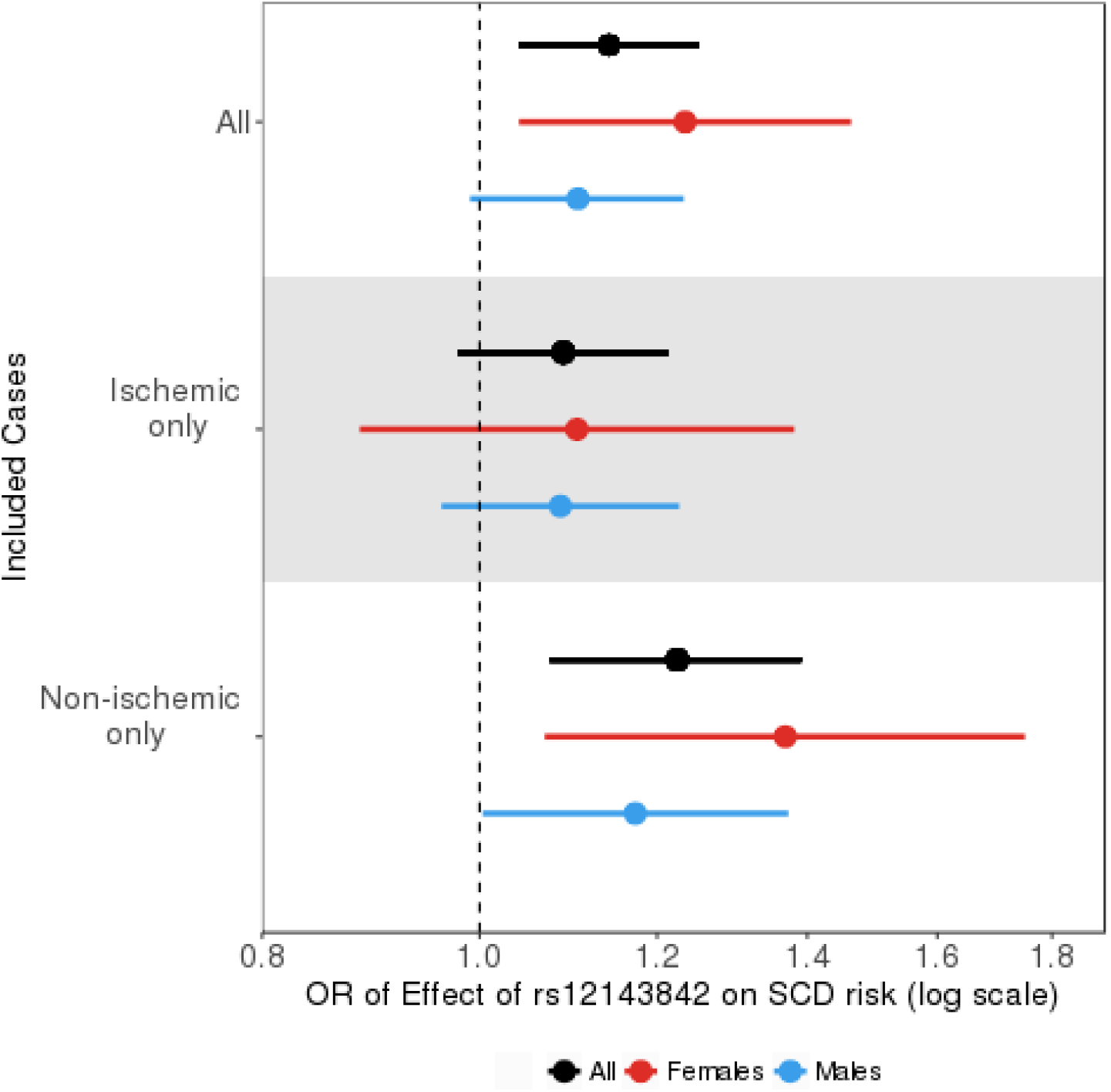
Forest plot of the association of rs12143842 with SCD risk. The top white panel represents the analysis including all SCD victims (2,282 cases); the middle gray panel includes ischemic-only SCD victims (1,478 cases); and the bottom white panel includes only non-ischemic SCD victims (750 cases). The dots represent the odds ratio of the rs12143842 QT prolonging allele on SCD risk and the lines represent the 95% confidence intervals. Both sexes (black), females only (red), and males only (blue). Additional information found in **Supplementary Table 4**.

### Ischemic vs. Non-ischemic

To explore whether the association of rs12143842 differs by underlying disease pathology, we stratified the SCD cases into those with (1) underlying ischemic heart disease (n=1,478), (2) non-ischemic heart disease (n=750), and (3) other pathologies (myocarditis, cardiac anomaly, and normal autopsy, n=54). The rs12143842 T allele had the strongest association in non-ischemic SCD individuals with an OR of 1.23 (95% CI, 1.07-1.39; *P*=0.003). A weaker non-significant association was observed in both ischemic SCD individuals (OR=1.09; 95% CI, 0.98-1.21; *P*=0.12), and those with other underlying conditions (OR = 1.11; 95% CI, 0.71-1.73; *P*=0.64).

### Men vs. Women

Given that QT interval is a stronger SCD risk factor in men than women, and rs12143842 has a larger effect on QT interval in women than in men,^20^ we next investigated whether the effect of rs12143842 on SCD risk differed between men and women. We limited sex-stratified analyses to SCD cases with underlying ischemic and non-ischemic pathology and excluded those with other underlying conditions due to the small sample size of those with other conditions.

Among 1,862 SCD male victims and 1,641 male controls, the rs12143842 QT prolonging allele was marginally associated with an increased risk of SCD (OR of 1.11; 95% CI, 0.99-1.23; *P*=0.07). When stratified by underlying disease pathology, the association was significant among non-ischemic SCD males (579 cases/1,641 controls) with an OR of 1.17 (95% CI, 1.00-1.37; *P*=0.045), while there was no statistically significant association in ischemic SCD males (1,245 cases/1,641 controls) for SCD risk (OR =1.09; 95% CI, 0.96-1.23; *P*=0.18; *P* for difference between ischemic/non-ischemic males=0.48).

Overall, among 420 female SCD cases and 1,920 female controls, the rs12143842 QT prolonging allele was associated with increased SCD risk (OR of 1.24; 95% CI, 1.04-1.46; *P*=0.015). Similar to findings among men, a stronger association was observed in the non-ischemic SCD women (171 cases/1,920 controls), with the rs12143842 T allele association with a 1.37-fold increased SCD risk (95% CI, 1.07-1.75; *P*=0.013) than among ischemic SCD women (233 cases/1,920 controls) (OR = 1.11 for each copy of the variant allele; 95% CI, 0.88-1.38; *P*=0.39; *P* for difference between ischemic/non-ischemic women=0.08).

### Mendelian Randomization of QT Interval

Using Mendelian randomization approaches, we have previously established that QT interval is causally associated with SCD.^11^ To investigate whether these causal associations differ based on sex and underlying disease, we calculated genetic risk score association (GRSA) estimates using the genome-wide significant SNPs from the most recent QT interval GWAS.^19^ Inverse-weighted (IVW) linear regression was performed to compare the effect of the SNP on QT interval to the effect of the SNP on SCD risk in the sex-stratified and underlying disease-stratified datasets. Results are summarized in **Figure 2** and **Supplementary Table 5**.

**Figure 2.**
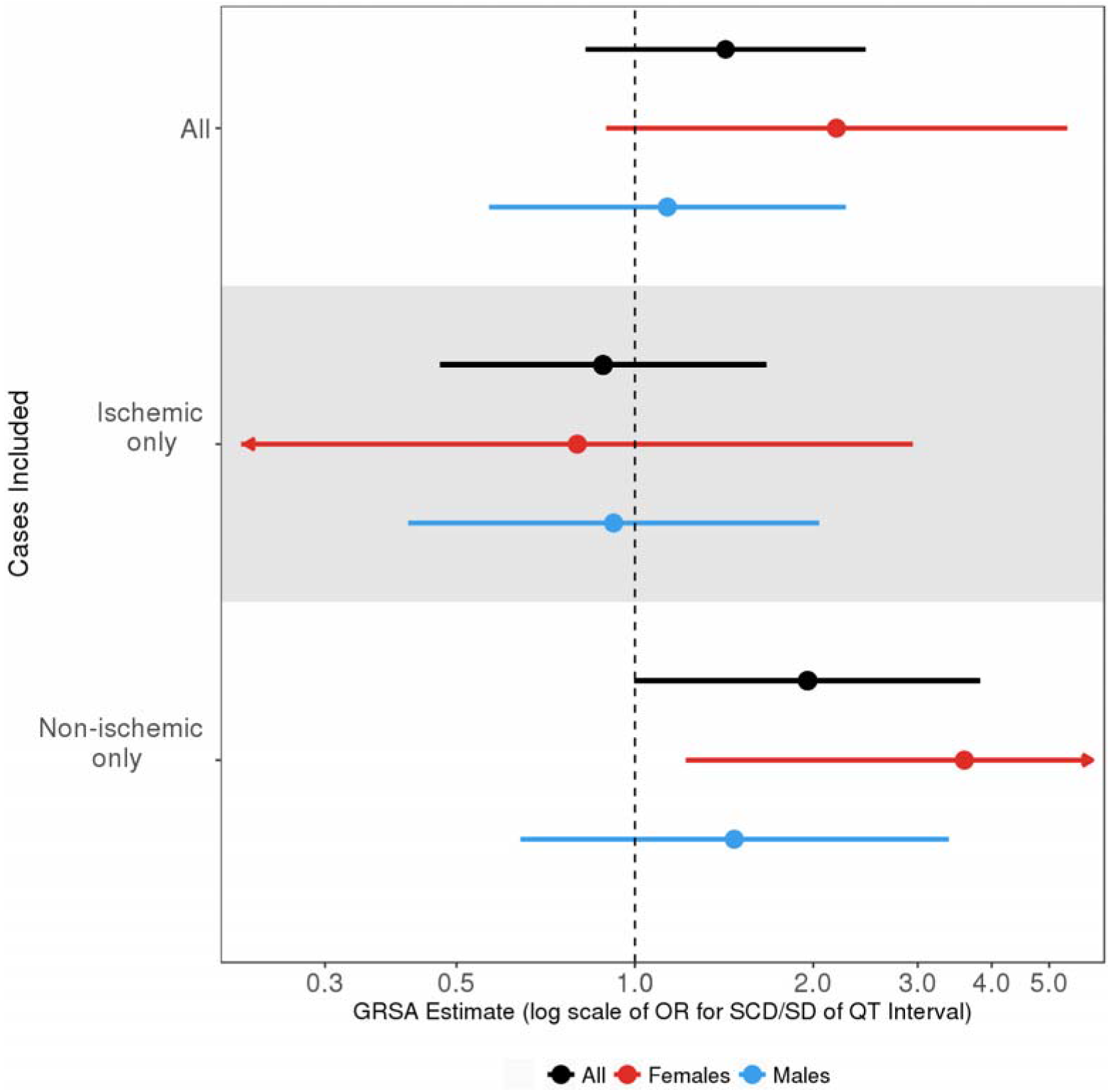
Genetic risk score association (GRSA) estimates for QT interval with SCD. The data points in the top plot represent the exponentiated GRSA estimates of QT interval on SCD (in log odds of SCD/SD of QT interval) and corresponding 95% confidence intervals. The top white panel represents the analysis including all SCD cases used in the MR analysis (1,168 cases); the middle gray panel includes ischemic-only SCD cases (611 cases); the bottom white panel includes only non-ischemic SCD cases (507 cases). Each panel includes analyses using: both sexes (black), females only (red), and males only (blue). Additional information found in **Supplementary Table 5**.

Among all SCD victims (n=1,168 cases/761 controls), a one standard deviation (SD) increase in QT interval was associated with a 1.42-fold increased risk of SCD (95% CI, 0.83-2.45; *P*=0.20). While not statistically significant, these findings are consistent with our previous work (previous findings: odds ratio in cardiac arrest risk per SD increase in QT, 1.44; 95% CI, 1.13-1.83; *P*=0.018)^11^. Similar to our findings with *NOS1AP* locus SNP rs12143842, we found that the causal relationship of QT interval and SCD differs between individuals with ischemic heart disease and individuals with non-ischemic disease. Among non-ischemic SCD victims (507 cases/761 controls), there was a 1.96-fold increase in SCD risk per SD increase in QT (95% CI, 1.00-3.82; *P*=0.05). By contrast, there was no evidence of a causal association of QT interval with SCD among SCD cases with ischemic disease (611 cases/761 controls; OR= 0.88; 95% CI, 0.47-1.67; *P*=0.70).

Non-ischemic female SCD cases had the strongest causal association of QT interval with SCD (odds ratio in SCD risk per SD increase in QT, 3.60; 95% CI, 1.22-10.59; *P*=0.02). Non-ischemic males had a large but non-significant causal association estimate between QT interval and SCD (odds ratio in SCD risk per SD increase in QT, 1.47; 95% CI, 0.64-3.39; *P*=0.36). Among those with underlying ischemic disease, there was no evidence for a causal relationship of QT interval with SCD for men or women (odds ratio in SCD risk per SD increase in QT, 0.92; 95% CI, 0.41-2.05; *P*=0.84 and odds ratio in SCD risk per SD increase in QT, 0.80; 95% CI 0.22-2.94; *P*=0.74, respectively).

## DISCUSSION

In the general population, women have longer QT intervals than men; women experience a higher rate of arrhythmias in the setting of prolonged QT interval; and prolonged QT interval is causally associated with SCD. We therefore hypothesized that women would show a greater association between genetically determined prolonged QT interval and SCD. Given the different etiologies between ischemic and non-ischemic cardiac disease, we further hypothesized that the genetic association with prolonged QT interval would also differ between the different underlying diseases. Our results, while not conclusive, support both of these hypotheses. Overall, we found that rs12143842, the top QT interval-associated SNP from previous GWAS^19^, was associated with SCD risk in our overall dataset. We observed a stronger association with SCD risk in non-ischemic individuals than ischemic individuals. Furthermore, women with SCD in the setting of non-ischemic cardiac disease had the strongest association between rs12143842 and SCD risk. Our Mendelian randomization analyses had similar findings; non-ischemic individuals showed a causal association between prolonged QT interval and SCD, and female non-ischemic individuals had the strongest causal association. By contrast, both the SNP association and Mendelian randomization analyses did not show evidence for a genetic (causal) association between QT interval and SCD due to underlying ischemic disease in men or women. These results suggest that SCD in the setting of ischemic disease may not be strongly influenced by myocardial repolarization (QT interval), or that the effect of prolonged QT interval on ischemic SCD risk is masked by other risk factors exerting a larger effect. While the differences in sex- and underlying disease-stratified associations were not statistically significant, these findings are nevertheless consistent with our underlying hypotheses; together these results provide evidence that SCD risk in non-ischemic individuals, particularly women with non-ischemic disease, is influenced by genetically determined QT interval.

The underlying cause(s) of the sex differences in the association between prolonged QT interval and SCD remains unknown, however, sex hormones may play a role. Studies have previously established that testosterone and progesterone shorten the QT interval, while estrogen lengthens the QT interval.^28,29^ While the underlying mechanism is unknown, our findings support the hypothesis that non-ischemic individuals are more susceptible to the effects of QT interval prolongation on developing SCD. Given that women already have underlying prolonged QT due to sex hormones, the addition of QT prolonging genetic susceptibility (i.e. the T allele of the *NOS1AP* SNP rs12143842) may result in the higher observed risk of SCD in women with non-ischemic disease.

While our study provides evidence for differences in SCD risk factors between both underlying disease and sex, several limitations should be noted. First, many of our analyses did not meet traditional statistical significance cut-offs, though we note that the results are entirely consistent with our original hypotheses. Thus, additional samples with autopsy-confirmed disease are necessary to confirm our results. Second, there is likely additional phenotypic heterogeneity within the underlying disease subgroups. The non-ischemic group, as noted in the supplementary methods, consists of eight different cardiac conditions. It is possible these different conditions, while similar in nature, may differ in their relationship between QT interval and SCD risk. Additional samples are needed to further stratify the non-ischemic group to investigate whether a particular condition is driving the association. Third, while our MDS components indicated potential population substructure within a subset of samples, when we included the components as covariates in our analysis, the effect was actually stronger. Therefore, not adjusting our main analysis for population substructure is likely resulting in a downward bias of the true association. Fourth, the NFBC1966 cohort used for our controls consisted of relatively young individuals (31 years old). Given the mean age of our SCD cohort was 60 years old, it is likely some of our “controls” will go on to have an SCD event later in life, and by not excluding these individuals, we bias our estimates towards the null. Lastly, the most interesting and strongest associations were seen in women and since women on average have lower rates of SCD, we have the least power to detect differences within this group. Nevertheless, our findings that female SCD victims with non-ischemic disease had the greatest association between prolonged QT interval and SCD risk were consistent between the various analyses performed, including both SNP association tests and Mendelian randomization. This is consistent with our original hypothesis, which stated that the effect of prolonged QT interval will differ by underlying disease pathology and would be stronger in females than males.

In conclusion, our study of autopsy-confirmed SCD victims provides evidence to support the hypotheses that SCD risk factors, specifically prolongation in QT interval, differ by both the underlying disease and sex. Non-ischemic SCD victims demonstrated a stronger genetic association, as well as a potentially causal association, between prolonged QT interval and SCD risk and these associations were strongest in female non-ischemic SCD individuals. SCD victims with underlying ischemic disease did not provide evidence for a strong genetic association, nor a causal association, between prolonged QT interval and SCD, regardless of sex. These findings provide additional evidence that SCD risk factors, particularly prolonged QT interval, differ between sex and underlying disease etiology.

## Data Accession

The data/analyses presented in the current publication are based on the use of study data downloaded from the dbGaP web site, under phs000276.v2.p1.

## Fingesture

This work was supported by the Juselius Foundation (Helsinki, Finland); the Council of Health of the Academy of Finland (Helsinki, Finland); the Montreal Heart Institute Foundation; Finnish Foundation for Cardiovascular Research (Helsinki, Finland); and Erkko Foundation (Helsinki Finland).

## NFBC1966

The NFBC1966 Study is conducted and supported by the National Heart, Lung, and Blood Institute (NHLBI) in collaboration with the Broad Institute, UCLA, University of Oulu, and the National Institute for Health and Welfare in Finland.

## ACKNOWLEDGEMENT

The authors thank all the staff and participants from the studies contributing to this manuscript for their important contributions.

## NFBC1966

This manuscript was not prepared in collaboration with investigators of the NFBC1966 Study and does not necessarily reflect the opinions or views of the NFBC1966 Study Investigators, Broad Institute, UCLA, University of Oulu, National Institute for Health and Welfare in Finland and the NHLBI.

