## Supplementary materials for "The effect of sex and underlying disease on the genetic association of QT interval and sudden cardiac death"

**SUPPLEMENTAL MATERIAL**

A.

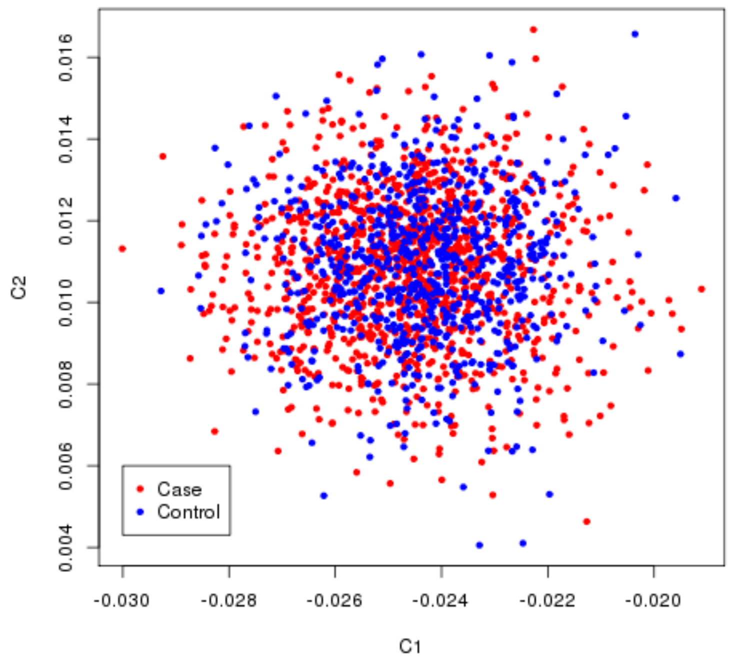


B.

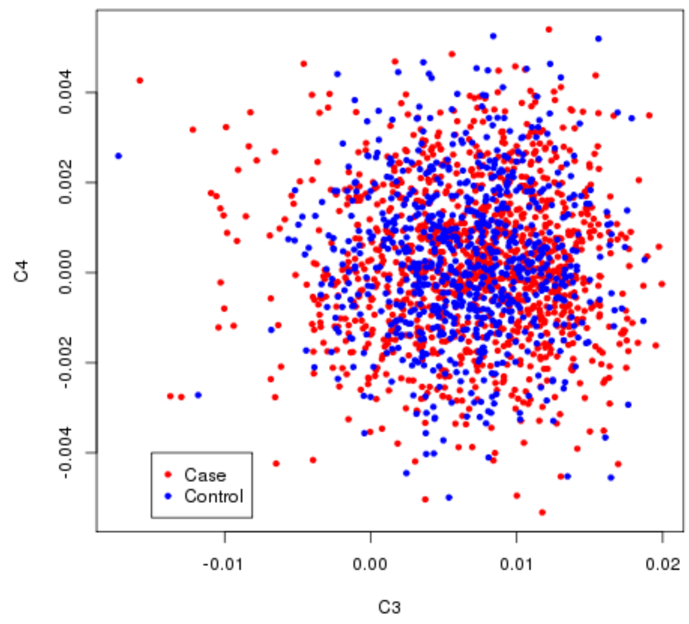

C.

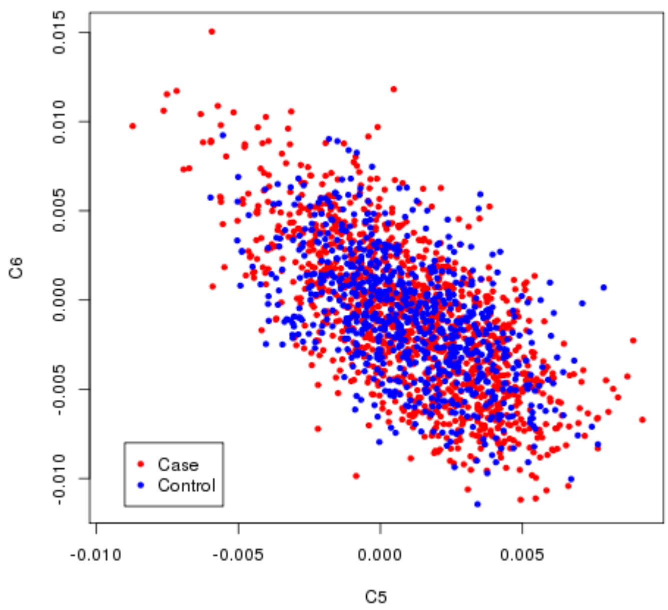
0.015
0.010
0.005
0.000
-0.005
-0.010
C6
Case
Control
-0.010
-0.005
0.000
0.005
C5

D.

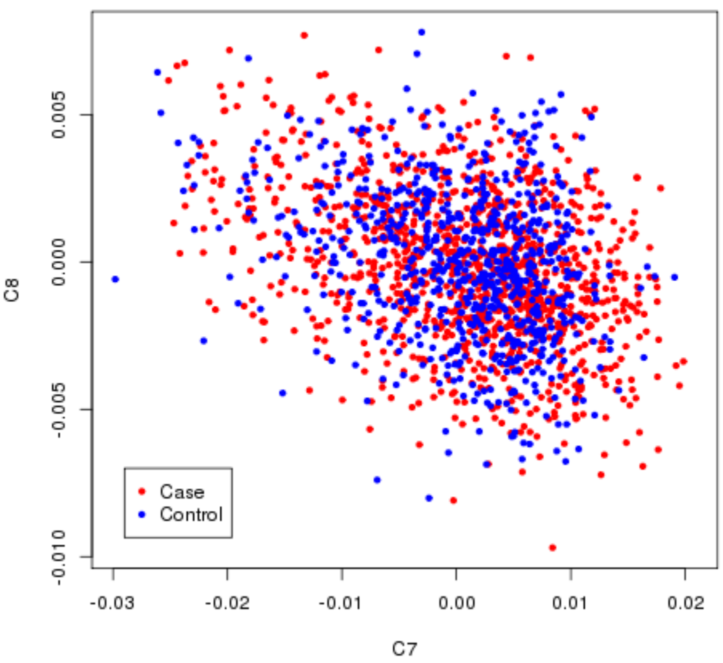
0.005
0.000
-0.005
-0.010
C8
Case
Control
-0.03
-0.02
-0.01
0.00
0.01
0.02
C7

E.

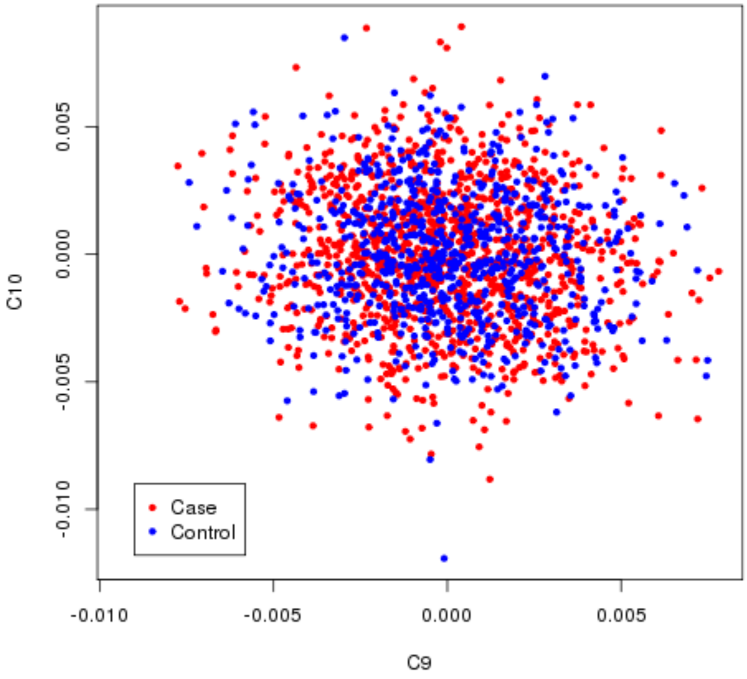


**Supplementary Figure 1. Multi-dimensional scaling (MDS) plot of Fingesture and NFBC1966 cohort samples.** The plots A-E demonstrates strong genetic overlap between the Fingesture cohort (red) and the NFBC1966 cohort (blue).

**Table S1. Genotyping Platform Sample Characteristics**

| Genotyping platform | Illumina Infinium Global Screening Array (GSA) | Affymetrix Genome-wide Human SNP Array 6.0 | Agena Biosciences MassARRAY | AB Taqman | Illumina Sequencing |
|---|---|---|---|---|---|
| N, number of total cases | 1168 | 358 | 574 | 572 | 825 |
| N, number of total controls | 761 | NA | 422 | 2175 | 563 |
| N, number of independent cases* | 1168 | 315 | 122 | 496 | 181 |
| N, number of independent controls* | 761 | NA | 251 | 2140 | 408 |
| QC critieria | Sample and SNP call rate (<95%); sex check; duplicate removal; cryptic relatedeness; genetic outlier removal using PCA | Sample and SNP call rate (<95%); sex check; duplicate removal; cryptic relatedeness; genetic outlier removal using PCA | Sample and SNP call rate (<95%) | NA | Minimum SNP read depth (10x); Sample and SNP call rate (<95%); sex check; duplicate removal; cryptic relatedeness; genetic outlier removal using PCA |
| Sex, number of women among independent cases | 218 | 50 | 30 | 91 | 31 |
| Sex, number of women among independent controls | 407 | NA | 145 | 1140 | 228 |
| Age, mean age at SCD event | 60.1 | 62.8 | 59.8 | 64.3 | 58 |
| N, number of ischemic SCD cases | 610 | 310 | 44 | 427 | 87 |
| N, number of non-ischemic SCD cases | 557 | 5 | 78 | 69 | 94 |
| Number of non-matching alleles between overlap samples† | 0 | 0 | 1 | 0 | 1 |

*This indicates the number of non-overlapping samples in each dataset used in the SNP analysis; overlapping samples were prioritized in the following order: (1) GSA; (2) Affy; (3) Sequencing; (4) MassARRAY; (5) Taqman

†Non-matching samples were removed from both datasets

**Table S2. Multi-dimensional scaling (MDS) regression results**

| Covariate | Beta | SE | *P* |
|---|---|---|---|
| Sex | -1.61 | 0.105 | <0.001 |
| MDS Component 1 | -0.296 | 0.328 | 0.36 |
| MDS Component 2 | -0.231 | 0.288 | 0.42 |
| MDS Component 3 | 0.477 | 0.189 | 0.011 |
| MDS Component 4 | -0.265 | 0.292 | 0.37 |
| MDS Component 5 | 0.099 | 0.269 | 0.71 |
| MDS Component 6 | 0.147 | 0.245 | 0.55 |
| MDS Component 7 | 0.316 | 0.102 | 0.002 |
| MDS Component 8 | 0.009 | 0.210 | 0.97 |
| MDS Component 9 | -0.002 | 0.197 | 0.99 |
| MDS Component 10 | 0.001 | 0.194 | 0.99 |

*Components were re-scaled by mulitiplying by 100 before regression to avoid numerical errors in R

**Table S3. Multi-dimensional scaling (MDS) regression results for rs12143842**

| Covariates used in model | Beta | SE | *P* | Variance Explained |
|---|---|---|---|---|
| Sex | 0.211 | 0.083 | 0.011 | 0.101 |
| Sex + MDS Components 1-10 | 0.227 | 0.084 | 0.007 | 0.108 |

**Table S4. rs12143842 SNP association results**

| | All | | | | |
|---|---|---|---|---|---|
| **Dataset** | **cases/controls** | **Beta** | **SE** | ***P*** | ***P* for ischemic/ non-ischemic difference** |
| All cases/population controls | 2282/3561 | 0.133 | 0.047 | 0.005 | |
| ischemic cases/population controls | 1478/3561 | 0.086 | 0.055 | 0.11 | 0.15 |
| non-ischemic cases/population controls | 750/3561 | 0.203 | 0.067 | 0.003 | |
| Other cases/population controls | 54/3561 | 0.106 | 0.226 | 0.64 | |

| | Males only | | | |
|---|---|---|---|---|
| **Dataset** | **cases/controls** | **Beta** | **SE** | ***P*** |
| All cases/population controls | 1862/1641 | 0.101 | 0.056 | 0.07 |
| ischemic cases/population controls | 1245/1641 | 0.083 | 0.062 | 0.18 |
| non-ischemic cases/population controls | 579/1641 | 0.160 | 0.080 | 0.045 |
| Other cases/population controls | 38/1641 | -0.172 | 0.290 | 0.55 |

| | Females only | | | | |
|---|---|---|---|---|---|
| **Dataset** | **cases/controls** | **Beta** | **SE** | ***P*** | ***P* for male/female difference** |
| All cases/population controls | 420/1920 | 0.211 | 0.087 | 0.015 | 0.29 |
| ischemic cases/population controls | 233/1920 | 0.100 | 0.114 | 0.39 | 0.79 |
| non-ischemic cases/population controls | 171/1920 | 0.314 | 0.126 | 0.013 | 0.14 |
| Other cases/population controls | 16/1920 | 0.649 | 0.377 | 0.09 | 0.015 |

**Table S5. Mendelian randomization of QT interval results**

| | All | | | | |
|---|---|---|---|---|---|
| **Dataset** | **cases/controls** | **SNPs included** | **GRSA Estimate [95% CI]** | ***P*** | ***P* for ishemic/non-ischemic difference** |
| All cases/population controls | 1168/761 | 57 | 0.352 [-0.191, 0.895] | 0.20 | |
| ischemic cases/population controls | 611/761 | 57 | -0.124 [-0.757, 0.510] | 0.70 | 0.09 |
| non-ischemic cases/population controls | 507/761 | 57 | 0.671 [-0.003, 1.340] | 0.05 | |

| | Males only | | | |
|---|---|---|---|---|
| **Dataset** | **cases/controls** | **SNPs included** | **GRSA Estimate [95% CI]** | ***P*** |
| All cases/population controls | 950/354 | 57 | 0.126 [-0.567, 0.820] | 0.72 |
| ischemic cases/population controls | 528/354 | 57 | -0.083 [-0.881,0.716] | 0.84 |
| non-ischemic cases/population controls | 387/354 | 57 | 0.386 [-0.445,1.220] | 0.36 |

| | Females only | | | | |
|---|---|---|---|---|---|
| **Dataset** | **cases/controls** | **SNPs included** | **GRSA Estimate [95% CI]** | ***P*** | ***P* for male/female difference** |
| All cases/population controls | 218/407 | 57 | 0.783 [-0.112, 1.680] | 0.09 | 0.26 |
| ischemic cases/population controls | 83/407 | 57 | -0.224 [-1.530, 1.080] | 0.74 | 0.86 |
| non-ischemic cases/population controls | 120/407 | 57 | 1.28 [0.202, 2.360] | 0.020 | 0.20 |
